# Structural characterization of covalently stabilized human cystatin C oligomers

**DOI:** 10.1101/654772

**Authors:** Magdalena Chrabąszczewska, Adam K. Sieradzan, Sylwia Rodziewicz-Motowidło, Anders Grubb, Christopher M. Dobson, Janet R. Kumita, Maciej Kozak

## Abstract

Human cystatin C (HCC), a cysteine-protease inhibitor, exists as a folded monomer under physiological conditions but has the ability to self-assemble via domain swapping into multimeric states, including oligomers with a doughnut-like structure. The structure of the monomeric HCC has been solved by X-ray crystallography, and a covalently linked version of HCC (stab-1 HCC) is able to form stable oligomeric species containing 10-12 monomeric subunits. We have performed molecular modeling, and in conjunction with experimental parameters obtained from AFM, TEM and SAXS measurements, we observe that the structures are essentially flat, with a height of about 2 nm, and the distance between the outer edge of the ring and the edge of the central cavity is ~5.1 nm. These dimensions correspond to the height and diameter of one stab-1 HCC subunit and we present a dodecamer model for stabilized cystatin C oligomers using molecular dynamics simulations and experimentally measured parameters. Given that oligomeric species in protein aggregation reactions are often transient and very highly heterogeneous, the structural information presented here on these isolated stab-1 HCC oligomers may provide useful to further explore the physiological relevance of different structural species of cystatin C in relationship to protein misfolding disease

## 1. Introduction

Human cystatin C (HCC), containing 120 amino acids, belongs to the cystatin type 2 superfamily[1, 2], and is a potent inhibitor of papain-like (C1) and legumain-like (C13) cysteine-proteases[3, 4]. In humans, HCC was originally identified in cerebrospinal fluid (CSF), but has subsequently been found in other bodily fluids and tissues[5–7]. The wild-type (WT) form of HCC is a component of amyloid deposits in, mostly elderly, patients with sporadic cerebral amyloid angiopathy[8]. Interestingly, the L68Q variant of HCC is associated with a rare hereditary cystatin C amyloid angiopathy, where the protein forms amyloid deposits in patients suffering from hereditary cerebral hemorrhage with amyloidosis[8–10]. To date, the crystal structures of monomeric and dimeric WT HCC have been characterized, the latter in two polymorphic forms[11–13], along with monomeric and dimeric crystal structures of several HCC mutational variants[14, 15]. Under physiological conditions WT HCC is a monomer, but when crystallized the protein readily forms domain-swapped dimers and when subjected to mildly destabilizing solvent conditions, it forms a number of oligomeric states and fibrils through this domain swapping phenomenon [12, 16]. An engineered variant of cystatin C (stab-1 HCC), which contains an additional disulfide bridge (L47C-G69C), stabilizes the monomeric form of the protein and reduces the ability of the protein to form fibrils[14, 17]. Stable oligomeric species of stab-1 HCC can be formed following incubation of high concentrations of monomeric protein at pH 7.4 in the presence of 1M guanidinium hydrochloride and the reducing agent, dithiothreitol[16]. These oligomers can be separated by size-exclusion chromatography (SEC) and are SDS-resistant but reducing agent sensitive, suggesting that intermolecular disulfide bonds stabilize the oligomers. By gel electrophoresis analysis, it is possible to isolate oligomers that have molecular weights which correspond to the size-range for decamers to dodecamers. These oligomer species do not retain their ability to inhibit papain-like cysteine proteases which indicates that the N-terminal loops 1 and 2 are not accessible, suggesting that these oligomers are formed via a domain-swapping mechanism[16].

Interestingly, the A11 antibody, designed to bind to soluble oligomers of the Aβ peptide, whose aggregation is associated with Alzheimer’s disease, is able to recognize a number of oligomeric species composed of different peptides and proteins[18], and has a weak affinity for the doughnut-like cystatin C oligomers[16]. This observation suggests that structural commonalities exist within different oligomeric species and therefore, detailed structural analysis of oligomers from an array of proteins is likely to prove to be insightful for understanding specific mechanisms of protein misfolding. At present, there are approximately 50 diseases linked to protein misfolding and amyloid deposition, including neurodegenerative disorders such as Alzheimer’s and Parkinson’s diseases, transmissible spongiform encephalopathies (TSEs), and non-neuropathogenic conditions such as systemic amyloidosis and type II diabetes[19, 20]. Although the primary and tertiary structures of the functional states of the peptides and proteins involved with these diseases are diverse, a hallmark of these disorders is the deposition of fibrillar structures which are remarkably similar[21].

The process of fibril formation involves a heterogeneous mixture of different aggregated species, including the mature fibrils, protofibrils and smaller, oligomeric species[22, 23]. Given that oligomer species are often transient, heterogeneous in nature and hard to isolate, many methods for producing stabilized oligomeric species have been employed with different protein substrates. These methods include the use of chemical crosslinking[24], altering ionic strength and buffer conditions[25], using lyophilization[26] and through chemical modifications[27]. These well-defined oligomer species have been used to search for structural attributes that may be important for pathogenicity, and increasing evidence has suggested that these oligomeric species are likely to be responsible for cellular toxicity through interactions and disruption of cellular membranes [18, 28–30]. Structural characterization of a range of such oligomer species have shown that they can adopt a number of morphologies, including spherical beads (2-5 nm in diameter), beaded chains, curly chains and ring-shaped (doughnut-like) structures[19, 20]. Though less commonly observed than other types of structures, doughnut-like oligomeric forms have been reported for human cystatin C (HCC)[31] and other amyloidogenic proteins, including α-synuclein, the amyloid β (Aβ) peptide and immunoglobulin light chains[32–34].

The aim of this present study is to define a structural model of the stable human cystatin C oligomers by combining information from the crystal structure of monomeric stab-1 HCC[14], along with experimental measurements of the oligomers obtained using transmission electron microscopy (TEM), atomic force microscopy (AFM) and small-angle X-ray scattering (SAXS) techniques, with molecular dynamics simulations. Using these techniques, we propose a dodecamer model of the stab-1 cystatin C oligomers.

## 2. Results

### 2.1. Overall morphology of the stab-1 HCC oligomers

TEM and AFM images were used to determine the overall geometric parameters of the stab-1 HCC oligomers that were isolated from size exclusion chromatography and correspond to 10-12 subunits. These samples provided the reference data for the construction of the initial molecular models of the oligomers. Representative TEM images, illustrating the morphology of the oligomers are shown in Figure 1. It is clear that a number of small, ring-like aggregates are present within the sample and that the predominant species appear to be approximately circular, having a diameter of 20-30 nm with a distinctive central cavity, in agreement with previous observations[31]. Some larger oligomers, of the order of 50 nm in diameter, and also the occasional short fibrillar species (Figure 1) can be observed, though much less frequently. Analysis of numerous TEM images at higher resolution (Figure 2) again reveals examples of circular, doughnut-shaped oligomers with differing sizes, with the smallest about 16-17 nm in diameter (d1) and the largest are about 20-24 nm. For the oligomers analyzed in detail, the distance between the edge of the central cavity and the inner edge of the outer ring (d2) is quite similar (~5-6 nm) (Figure 2); this value corresponds very well to the cross-sectional diameter of one cystatin C molecule in the crystal structure [14].

**Figure 1.**
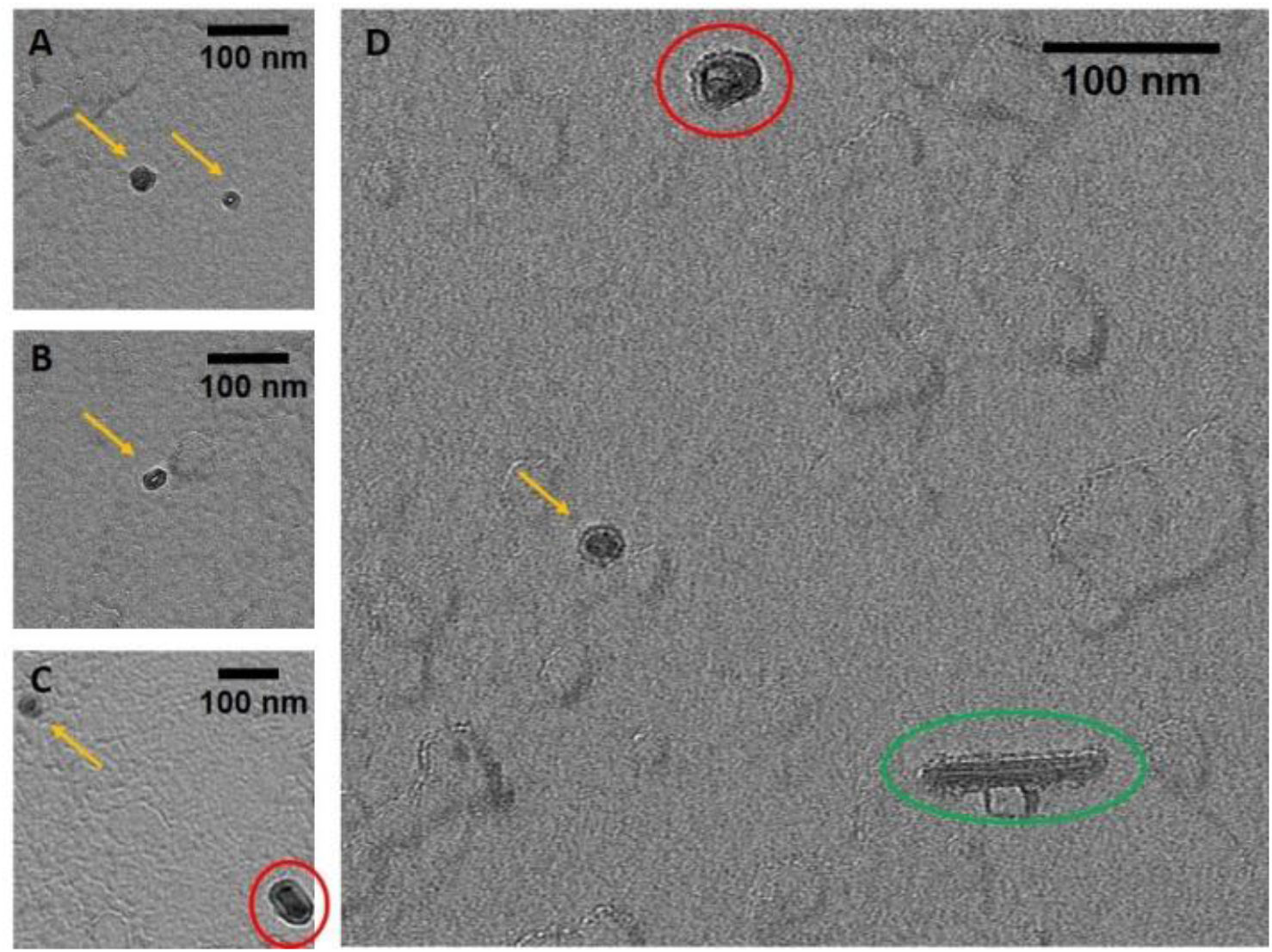
TEM images of the stab-1 HCC oligomers. Selected TEM images of the stab-1 HCC doughnut-like oligomers stained with uranyl acetate (yellow arrows, panels A-D) showing representative species present in the samples. Small numbers of short fibrils (green oval, panel D) and higher molecular weight oligomers (red circles, panels C, D) are also evident. Scale bars are 100 nm in each case.

**Figure 2.**
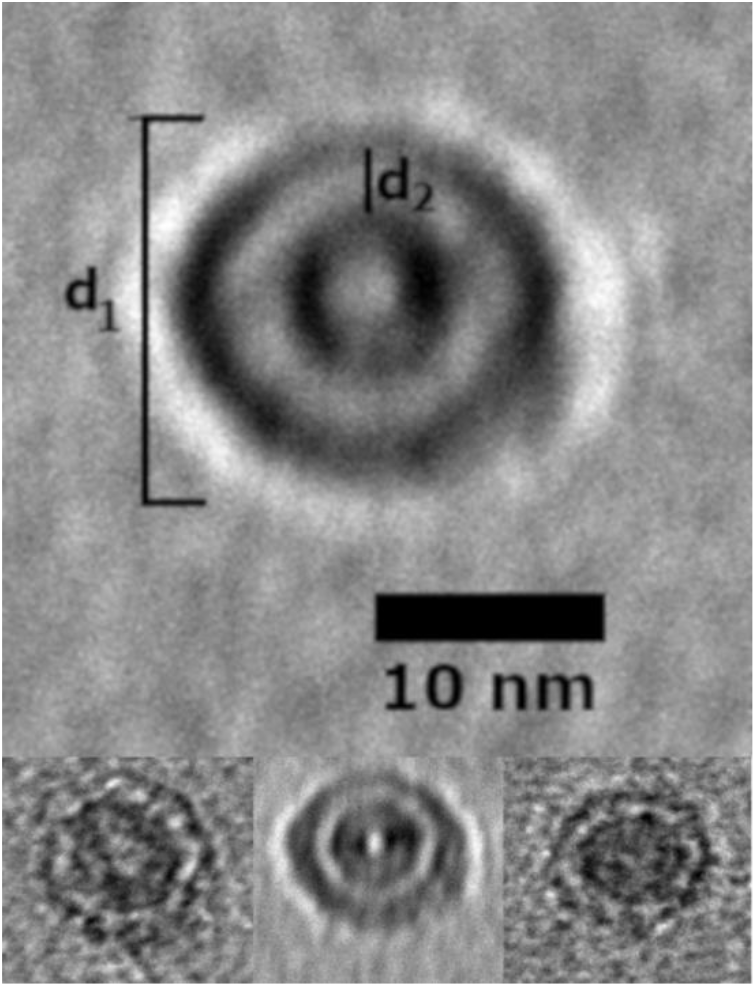
Selected stab-1 HCC doughnut-like images of oligomers obtained from TEM data. Top, enlargement of the images of stab-1 HCC oligomers observed by TEM. The diameter is indicated by “d1” and the distance between the inner and outer rings defined as “d2”. Bottom, variations in d1 and d2 are likely to be due to the differences in the electron density of the contrasting uranyl acetate adsorbed to the oligomer surface. Scale bars are 10 nm in each case.

To complement the TEM analysis, we used atomic force microscopy (AFM) to provide independent evidence about the overall morphology of the stab-1 HCC oligomers and also to determine the heights of the individual oligomers. Representative AFM images are shown in Figure 3, with the predominant species displaying the spherical morphology observed in the majority of the TEM images. The cross-sectional profiles (Figure 3, right panel), indicate that these circular oligomers have heights in the range of 1.6 to 2.5 nm and diameters of 20-30 nm which is slightly larger than what was observed by TEM. However, the AFM measurements were not calibrated to give accurate XY plane measurements in these experiments, and therefore these diameters may be over-estimated. The height measurements of the oligomers were therefore taken from the AFM studies, and the values for the diameters were based on the TEM measurements.

**Figure 3.**
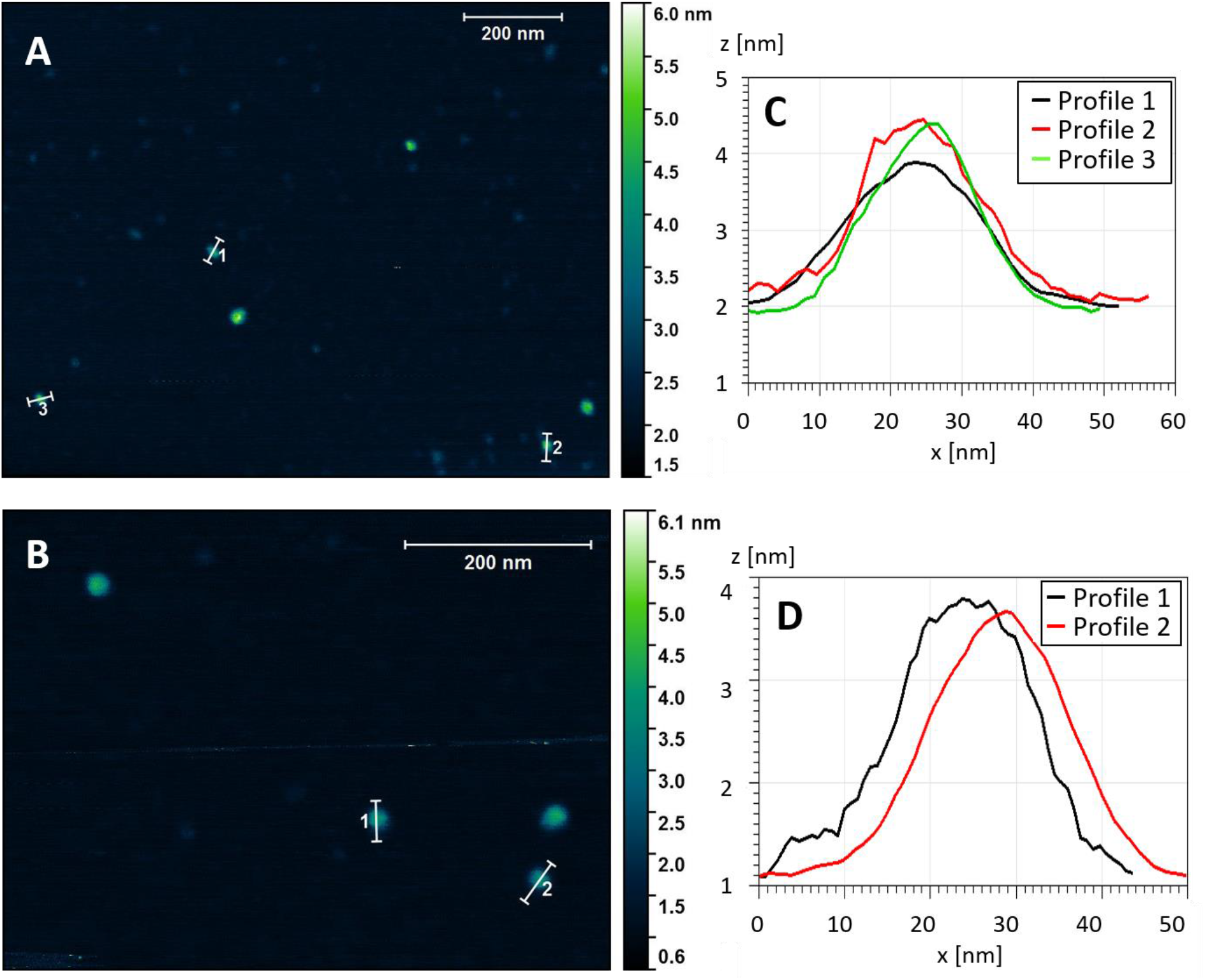
Topography of the stab-1 HCC oligomers. Selected AFM images of the stab-1 HCC oligomers (A, B) and cross-sectional profiles of the indicated oligomers in the images (C, D). The height profiles (z [nm]) are presented for the indicated oligomers from panel A (1-3) and panel B (1-2). From this analysis the diameter (x [nm]) can be approximated.

The crystal structure of the stab-1 HCC monomer shows that the protein molecule has a diameter of approximately 2.5-3.0 nm (excluding the hydration shell on the protein surface) (PDB code: 3GAX)[14]. Despite the possibility of errors in the heights measured by AFM as a result of strong adhesion to the mica surface, the value of 2 nm corresponds to a single layer of stab-1 HCC subunits.

### 2.2. Structural parameters of the stab-1 HCC oligomers in solution

SAXS studies of the stab-1 HCC oligomers in solution involved the use of synchrotron radiation which has a risk of causing the reduction of the disulfide bridges that stabilize the overall structure of the HCC subunits and also the oligomers. We know that human cystatin C and its variants are sensitive towards synchrotron radiation and prone to aggregation[35]. A detailed inspection of each of the recorded frames was therefore carried out and only those frames that did not show an evidence of radiolysis were selected for further analysis. The radius of gyration (R_g_) for the oligomers was determined by fitting the SAXS data (Supplementary Figure S1) to the Guinier equation. The result of the analysis gives a value for R_g_ of 5.28 ± 0.13 nm.

As a consequence of the limited s-range of the SAXS data due to the low available protein concentrations and some size polydispersity, a degree of heterogeneity is likely to exist. For these reasons, we were not able to conduct *ab initio* modeling but instead defined the geometric parameters of the flattened doughnut-like structure, which best depicts the stab-1 HCC oligomers using the universal R_g_ calculator[36]. These calculations indicate that the outer diameter of the doughnut-like structure is 20-23 nm, the height is 2.4-2.6 nm and the inner diameter is 7-8 nm. These values are consistent with the height and diameter parameters determined for a single stab-1HCC subunit of 16-24 nm (outer diameter) and 1.6-2.5 nm (height), obtained from TEM and AFM measurements. This information was therefore used to define reference parameters for the construction of the initial models of the stab-1 HCC doughnut-like oligomers, which were required as input into to molecular dynamics simulations (Table 1).

**Table 1.** Summary of structural parameters of the stab-1 HCC oligomers obtained by different experimental methods.

|  | Outer diameter (nm) | Inner diameter (nm)* | Height (nm) |
| --- | --- | --- | --- |
| TEM | 16-24 | 5-6 | - |
| AFM | 20-30 | - | 1.6-2.5 |
| SAXS | 20-23 | 7-8 | 2.4-2.6 |

\*The inner diameter is defined as the distance between the outer edge of the internal cavity and the outer edge of the whole ring structure.

### 2.3. Preliminary models of the stab-1 HCC oligomers

Two subtypes of stab-1 HCC oligomers, the decamers and dodecamers, were selected for molecular modeling on the basis of the geometric parameters, derived from TEM, AFM and SAXS data (Table 1), and also on criteria based on previously published biochemical studies [16]. It has been shown previously that the stab-1 HCC oligomers are not capable of inhibiting the protease papain, indicative of the papain-binding site (N-terminal loops 1 and 2) being buried within the oligomer structure, whereas the oligomers are still capable of inhibiting the activity of legumain, confirming that this part of the cystatin C structure is still available for protease-binding[16]. As these earlier studies strongly suggest that the domain-swapped dimeric structure is maintained, this was incorporated after selection from the initial modeling. For both the decamer and dodecamer oligomers, a geometric alignment of the HCC subunits using the stab-1 HCC monomeric crystal structure (PDB: 3GAX) and the SymmDoc platform[37] was performed. From a pool of 100 potential structures for each oligomer type obtained in this way, the models were grouped manually into five subfamilies (containing 2-12 models) differing to a small degree in the relative positions of the subunits.

Some of the models selected did not match the geometric parameters defined by the TEM, AFM and SAXS experimental data and these models were not taken forward into the molecular dynamics simulations; this eliminated a significant number of potential structures that contained inconsistent orientations of monomer-like subunits and also barrel-like arrangements that did not match the experimental dimensions. The models matching the experimental criteria within the decamer and the dodecamer subgroups were manually compared and four representative models from each of these sets were selected for future analysis. At this stage, three models of the stab-1 HCC decamers were selected for further MD simulations as the overall arrangement of subunits was different in each after the initial docking procedures, interestingly, one of these models was very unstable and lost its secondary and tertiary structures during the initial steps of the MD simulations; it was therefore excluded from further studies. A similar approach was taken for the dodecamer models, where just one dodecamer model was selected, which consisted of a very similar arrangement of subunits to the selected decamer models. All the initial models of stab-1 HCC decamers and dodecamers are shown in Supplementary Figure S2.

After selection of the starting models obtained from SymmDock, the fragment of the polypeptide chain (Pro78-Asn79) that is missing in the 3GAX structure was built using Swiss-PdbViewer v4.1[38], and a domain exchange between neighbouring units in the oligomers was implemented as shown in Supplementary Figure S3. This exchange was based on the domain swap mechanism observed in the native HCC crystal structures, in which the N-terminal fragment of the polypeptide chain (residues 1-58) was transferred from the first stab-1 HCC subunit to the second one, with the rest of the chain remained unaltered[11–13]. With these additions, the models were used for the molecular dynamics simulation step in AMBER.

### 2.4. Molecular dynamic simulations of the stab-1 HCC oligomers

After selecting the preliminary models as described above, we examined the structural and conformational stabilities of these stab-1 HCC oligomers using molecular dynamics simulations using the AMBER program package and the AMBER force field. To distinguish between the two decamer models, we denote them as stab-1 HCC decamer Model 1 and stab-1 HCC decamer Model 2. Molecular dynamics simulations of the stab-1 HCC decamer Model 1 and Model 2 and the dodecamer model were performed for a total of 50 ns per model. The changes in the potential energy of these systems as a function of simulation time are shown in Supplementary Figure S4. Throughout the MD simulations, the dodecamer model has a lower potential energy value compared to the models of the stab-1 HCC decamers. This is indicative of a greater stability of this dodecamer oligomer model relative to the decamer models and suggests that it is the most energetically favoured structure. After ca. 35 ns of MD simulations, the subsequent changes in energy for all models is relatively small even immediately after the reduction of the positional constraints imposed on Cα, indicating the stabilization of the energy parameters for the decamer and dodecamer structures.

For all the models studied here we also analyzed the changes in root mean squared deviations (RMSD) during the simulations. The changes in the positions of Cα atoms within all the configurations were compared to the coordinates in the initial model after minimization of the energies in solution. The changes in RMSD for all three models are shown in Supplementary Figure S4b. The rapid increase in RMSD values within the first 0.5 ns is likely to result from (i) bringing the system to the measurement temperature, (ii) the necessity for the protein structure to adapt to the in silico domain exchange, and (iii) the necessity of the initial structures to adapt after the transfer of the model, based on the protein structure from the crystallographic data environment, into solution by applying an *implicit* solvent model.

The potential energy and RMSD plots possess a step-like characteristic as a result of the stepwise reduction of the strength of the restraints imposed on the Cα atoms. Over the entire MD simulations, the change in RMSD values for the decamer Model 1, Model 2 and the dodecamer were 3.84 Å, 4.05 Å and 4.35 Å, respectively. The observed changes in RMSD are related to the presence of a number of conformational transformations within the native state of the protein in solution[39], and the larger initial conformational changes are likely to be due to local reorganization resulting from the domain exchange incorporated in the initial models. In all cases, the trajectory of the RMSD values stabilized throughout the time course of the MD simulations.

To complement these molecular dynamics simulations (time scale 50 ns), we performed canonical molecular dynamics using a scale-consistent UNitedRESidue (UNRES) course-grained force field [40–42]. This allowed us to do the simulations on a much longer time-scale (3 μs) and we report the RMSD trajectories along with the TM-scores, which are the scoring function for automated assessment of protein structure template quality. TM-scores allow evaluation of structural predictions which has no bias due to the target protein’s length or size and which uses all the residues of the modeled proteins in the evaluation of the score[43]. Simulations performed at 277K and 300K are shown in Figure S5 and Figure S6, and we found that after 3 μs at 277K the TM-scores were 0.52, 0.46 and 0.47 and at 300K, all three models exhibit similar TM-scores of 0.46, 0.43 and 0.43, for the dodecamer, decamer Model 1 and decamer Model 2, respectively. Therefore, as with our shorter-scale simulations, the most stable model was the dodecamer and we note that out of the 288 trajectories performed, no dissociation was observed (see comparison of final structures in Supplementary Figures S7 and S8). We further confirmed, using longer all-atom simulations, without any restraints, in explicit solvent, that decamer Model 1 and dodecamer are stable (Supplementary Figure S9). The RMSD for decamer Model 1 rose during the simulation to 7Å while the RMSD for the dodecamer rose to 16 Å. The lower value for decamer Model 1 revealed a higher stability for this structure as it remained virtually unchanged during the simulations (Supplementary Figure S10a), whereas the dodecamer shows signs of bending over the time course (Supplementary Figure S10b). Decamer Model 2 was found to be unstable during the all-atom simulations. Despite the fact that order of stability order for the decamer Model1 and the dodecamer differs between the all-atom simulations and the coarse-grained simulations, both methods confirm that there is no dissociation with either of these two models.

After the simulations, it was clear that the increase in RMSD is largely related to changes in the secondary structure within the oligomer models, as the tertiary structure and the overall shape, size and position of the stab-1 HCC monomer-like subunits within the oligomers do not change significantly (Figure 4a). For all the models of the stab-1 HCC oligomers, a partial loss of secondary structure is particularly evident within the β2 sheet and the L1 loop, which participates in domain swapping (Figure 4b). Thus, the structure for the entire subunit changes after domain swapping and results in the formation of a loop in place of the β sheet structure of the native state. A comparison of the secondary structure content within the final oligomer models to that of the stab-1 HCC monomer (PDB code: 3GAX) and the HCC dimer (PDB code: 1G96) is shown in Table 2. The highest degree of secondary structure within the polypeptide chain is present in decamer Model 1, with decamer Model 2 having the largest disruption to the secondary structure elements relative to the native monomer and dimer, containing a random coil fraction of 43%[12, 31].

**Figure 4.**
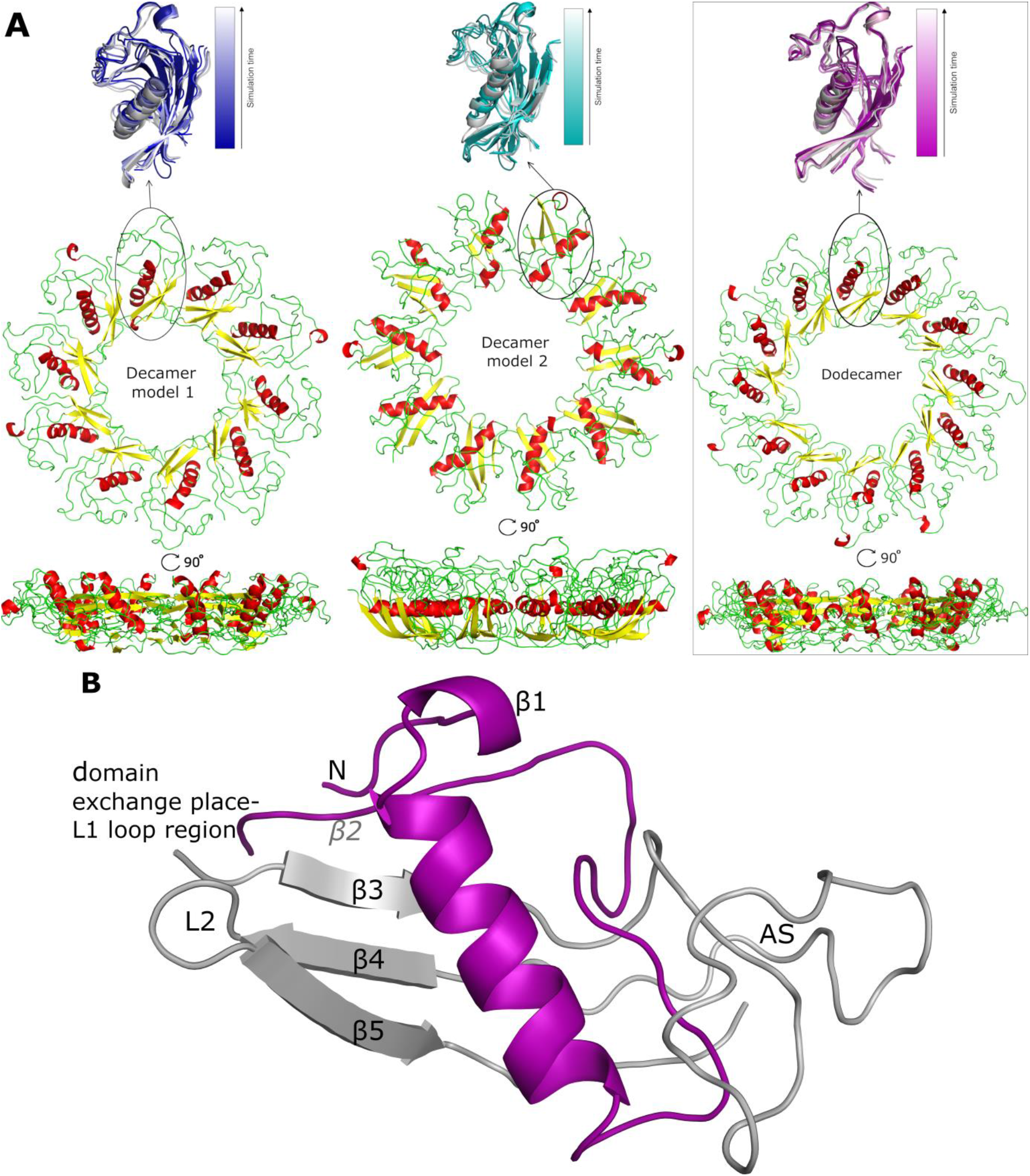
Models of the stab-1 HCC oligomers after MD simulations. A) Final models of HCC: decamers (left – Model 1, center – Model 2) and dodecamer (right) obtained in the molecular dynamics simulations. An enlargement of one subunit within the initial models (color) is superimposed onto the final structures (grey). B) Definition of secondary structure elements presented in the domain swapping region within the crystallographic model (molecule 1 - light gray, molecule 2 - purple). The appending structure (AS) is the broad random-coil region between strands β3 and β4 [12].

**Table 2.**
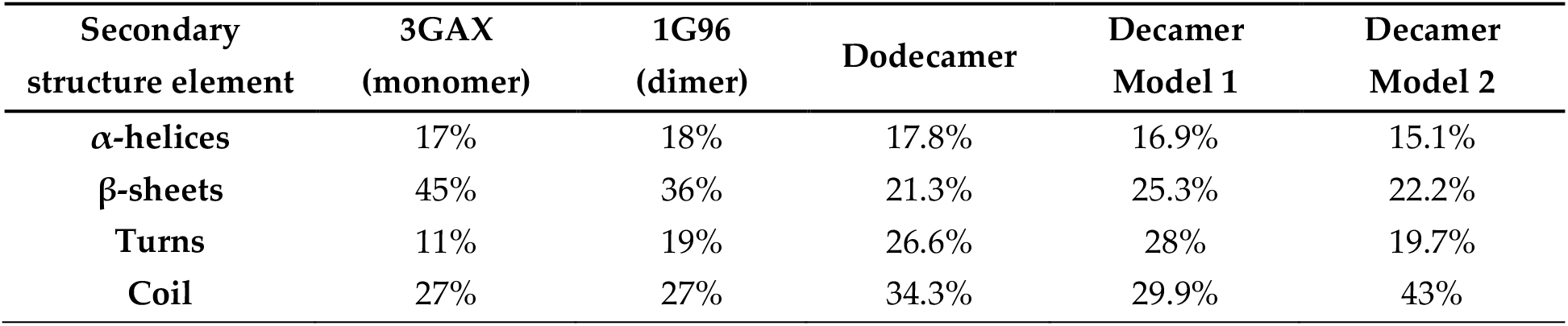
Secondary structure content of the proposed stab-1 HCC oligomer structures calculated using the STRIDE web server[44] and compared with values derived from the crystal structure of the stab-1 HCC monomer (PDB code: 3GAX;[14]) and the HCC dimer (PDB code: 1G96;[12]).

| Secondary structure element | 3GAX (monomer) | 1G96 (dimer) | Dodecamer | Decamer Model 1 | Decamer Model 2 |
| --- | --- | --- | --- | --- | --- |
| α-helices | 17% | 18% | 17.8% | 16.9% | 15.1% |
| β-sheets | 45% | 36% | 21.3% | 25.3% | 22.2% |
| Turns | 11% | 19% | 26.6% | 28% | 19.7% |
| Coil | 27% | 27% | 34.3% | 29.9% | 43% |

Analysis of the structures of the individual oligomer models show that the greatest structural changes occur within decamer Model 2 (see Table 2 and 3). In comparison to the initial model used for the MD simulations, the outer edge of this model has shifted to the interior of the oligomer structure, creating a bowl shape. The height of this oligomer also increases to 3.5 nm, a value that is higher than estimated from AFM measurements (1.6-2.4 nm). In contrast, the diameter of decamer Model 2 is 0.5 nm smaller than that observed for the decamer Model 1. Overall, the HCC dodecamer is the most energetically stable structure in the MD simulations and the dimensions of the proposed model are in closest agreement with the structural parameters of the circular oligomers obtained from experimental studies (TEM, AFM and SAXS) (Table 3).

**Table 3.** Structural parameters of models of the stab-1 HCC oligomers after MD simulations.

|  | Decamer Model 1 | Decamer Model 2 | Dodecamer |
| --- | --- | --- | --- |
| Outer Diameter [nm] | 12.0 | 11.5 | 13.8 |
| Inner Diameter* [nm] | 4.8 | 5.1 | 4.8 |
| Height [nm] | 2.3 | 3.5 | 2.5 |

\*Inner diameter is defined as the distance between the internal cavity and the outer ring structure

The conformational stability of the oligomers and their secondary structure depends on the presence of arrays of hydrogen bond networks that not only stabilize the structure of the particular subunits, but which are also observed to form between different subunits via domain swapping; a key factor for the stability of the entire oligomeric structure. For the dodecamer model, stable hydrogen bonds between two domains within the same subunit are formed between Asp65-Arg51 (O^D2^-H^H22^-N^H2^), and are present for 78.4% of the MD simulation time; domain swapped hydrogen bonds are, however, formed between two pairs of residues: Glu21-Arg51 (O^E1^-H^H11^-N^H1^) and Pro105-Asn61 (O-H^D22^-N^D2^) and are present for 84% and 72% of the MD simulations, respectively (Figure 5). The presence of domain swapped hydrogen bonds is indicative of the high stability of the dodecamer model.

**Figure 5.**
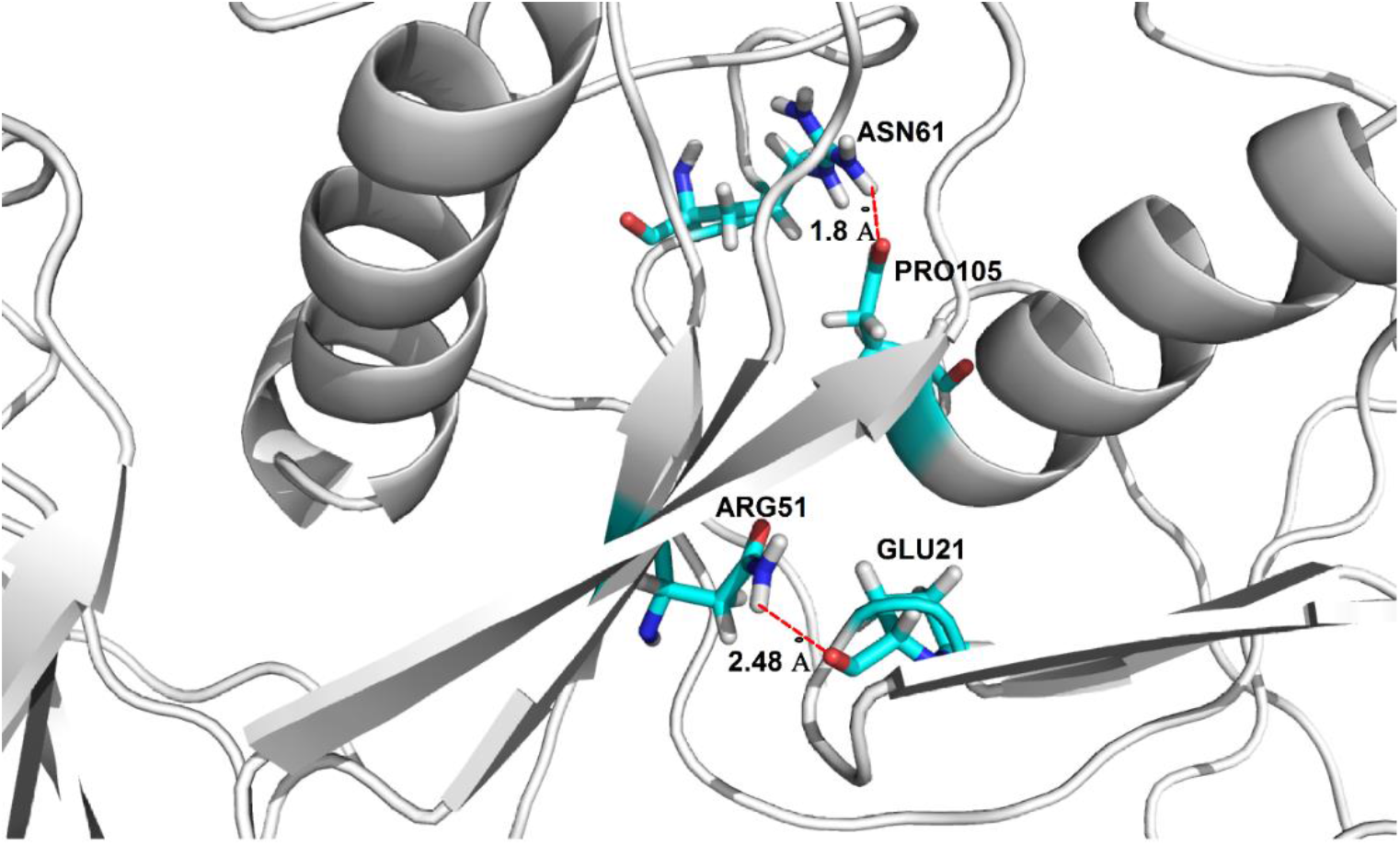
Location of domain swapping hydrogen bonds stabilizing the stab-1 HCC dodecamer oligomers. Domain swapping hydrogen bonds observed between Glu21-Arg51 and Pro105-Asn61.

## 3. Discussion

As a result of the good correlation between the MD simulations and the experimental parameters measured by AFM, TEM and SAXS, we propose a model of the stab-1 HCC oligomers in solution as a doughnut-like dodecamer structure. This dodecamer structure is the most energetically favoured of the three models we have examined. Although a small population of larger, cyclic species, as well as short fibrils, have been observed sporadically in TEM images, the majority of the species we have observed experimentally fit well with this proposed dodecamer model.

It is interesting to note that these stabilized HCC oligomers, which do not readily form fibrillar species, maintain a significant degree of native-like structure. Such maintenance of native-like structure has also been reported for WT HCC oligomeric species formed under non-denaturing conditions[45]. These reported oligomers retain native secondary structure and both papain and legumain enzymatic inhibitory function and appear to be non-domain swapped species, but unlike our stabilized oligomers, the average size of those species is that of a trimer[45]. Interestingly, stabilized oligomers from another member of the cystatin superfamily, stefin B (or cystatin B), have been observed and characterised as domain-swapped tetrameric oligomers which are stabilized by a “hand-shake mechanism” [46]. As such, Pro74 is in the *cis* form (opposed to the *trans* form normally adopted in the monomer) and this causes domain-swapped dimers to become intertwined. Like the stab-1 HCC oligomers, the reported trimeric non-domain swapped oligomers as well as the stefin B oligomers do not readily proceed to fibril formation of cystatin C[16, 41, 45, 46].

The stab-1 HCC aggregates analyzed in this study can be isolated and are relatively stable in solution, and this has enabled the detailed study of their structural nature. Using experimental measurements and MD simulations, we propose a dodecamer structural model of the stab-1 HCC doughnut-like oligomers. Given the generic property of proteins to form amyloid fibrils, mounting evidence that the formation of oligomers is also a commonality amongst diverse substrates is increasing and interestingly, along with oligomers isolated from intrinsically disordered proteins, such as Aβ peptides and α-synuclein, a number of structural studies have been reported for oligomers formed by proteins which have globular native structures, such as hen egg-white lysozyme, transthyretin, HypFN, acylphosphatase and cystatin C[47–50]. With the growing importance of understanding the biological significance of oligomeric species, the structural information presented here on the stab-1 HCC oligomers provides further information on the nature of such species.

## 4. Materials and Methods

### 4.1. Production of stabilized oligomers

Covalently stabilized oligomers of human cystatin C (stab-1 HCC) were produced and purified as described previously [16] and the samples for transmission electron microscopy and atomic force microscopy measurements were isolated by size-exclusion gel chromatography as previously detailed, and kept in sodium bicarbonate buffer (50 mM, pH 7.8, 4°C) prior to imaging.

### 4.2. Transmission electron microscopy

Stab-1 HCC oligomer solutions (3-5 μL) for characterization by TEM were applied to carbon-coated 400 mesh nickel grids (TAAB Laboratory Equipment Ltd., Aldermaston, UK), allowed to adsorb for 60 s and then blotted using filter paper. Samples were stained with 2% (w/v) uranyl acetate for 30 s, and washed with deionized water. The samples were imaged on a FEI Tecnai G2 transmission electron microscope at the Cambridge Advanced Imaging Centre (CAIC), University of Cambridge, UK. Images were recorded using the SIS Megaview II Image Capture System (Olympus, Tokyo, Japan). Analysis of the resulting TEM images was performed using ImageJ [51].

### 4.3. Atomic force microscopy

Topographic images of the oligomers were collected using a NanoWizard AFM system (JPK Instruments AG, Berlin, Germany). Purified samples were first diluted in deionized water by a factor of 1000-5000, to give concentrations of ~2-10 pM. These solutions (5-10 μl) were placed on freshly prepared mica surfaces, adsorbed for 10-20 min, gently washed with small amounts of deionized water and dried using nitrogen gas. AFM imaging was performed in the intermittent (air) contact mode using a silicon nitride cantilever. The analysis of the images was carried out using Gwyddion 2.45 software [52].

### 4.4. Small angle X-ray scattering

The small angle X-ray scattering (SAXS) data for solutions of stab-1 HCC oligomers (1.1 mg/mL) were collected on the BioSAXS X33 bending magnet beamline, operated by EMBL [53, 54] at the DORIS III storage ring of DESY (Hamburg, Germany). The experiments were conducted in a standard manner using an autosampler, a hybrid photon counting detector (Pilatus 1M-W, Dectris, Baden-Daettwil, Switzerland) and synchrotron radiation 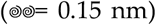. The data were processed using PRIMUS [55] from the ATSAS package [56], and the radius of gyration (*R_g_*) was calculated by fitting of the SAXS data in the *s*-range from 0.124 to 0.247 nm^−1^ 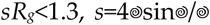, where 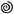 is a scattering angle) to the Guinier equation (Eq. 1)

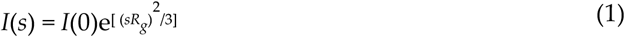

The universal *R_g_* calculator [41], was used to define the geometric parameters of the images that best depicted the shape of doughnut-like oligomers.

### 4.5. Molecular dynamics simulations

#### 4.5.1. Preparation of initial models

Preliminary models of stab-1 HCC decamers and dodecamers were constructed using the crystal structure of the stab-1 HCC monomer (PDB: 3GAX) [14]. The subunits (HCC monomers) were assembled into oligomers using the docking server SymmDock[37]. For the construction of the initial atomic models of the oligomers, two types of oligomers containing 10 and 12 monomer-like subunits were defined on the basis of previous gel filtration analysis [16], and the structures of the oligomers were obtained by the geometrical matching of the given subunits in space. The parameters (oligomer diameter, height and inner cavity diameter) determined from the TEM and AFM measurements were used as the initial geometric parameters. These dimensions suggest that the rings are composed of individual monomer-like subunits arranged in a circular shape. From the final set of 100 structures for each oligomer type produced using SymmDock, only those models satisfying the experimental geometrical and structural conditions were considered further. Once these models were selected, a fragment of the polypeptide chain (Pro78-Asn79) for which electron density is missing in the X-ray structure (3GAX) was built using Swiss-pdb Viewer 4.1, and the exchange of the subdomains, up to position Ala58, was implemented manually according to the domain swapping mechanism observed in the native HCC crystal structure [11–13]. The domain swapping is implemented intermolecularly throughout the oligomer therefore, we manually broke the bond between residue 58 and 59 to allow the transfer of residues 1-58 from one subunit to the next. Due to this broken bonds in these initial models, larger positional restraints were imparted at the beginning of the MD simulations. The atomic positions of the residues in the N-terminal segment of the polypeptide chain (residues 1-58) were transferred from the first stab-1 HCC monomer-like subunit to the second one, with the rest of the first chain remaining unaltered. This phase of the modeling was conducted in JOE (operating in the Linux environment). Analogous procedures were performed for all subunits in the selected models of the stab-1 HCC decamers and dodecamers prior to the molecular dynamics simulations in AMBER.

#### 4.5.2. Simulations using AMBER

Molecular dynamics simulations of the doughnut-like oligomers (decamers and dodecamers) were carried out using the PMEMD module of the program SANDER from the AMBER 12 package [57, 58], employing the graphics processing units (GPUs) [58]. Preliminary minimizations of the initial oligomeric structures were performed in two steps, the first involving minimization *in vacuo* and the second in *implicit* solvent. Each minimization cycle was completed within 10,000 steps, where each step was 2 fs in duration, and both the minimization steps and the molecular dynamics simulations were conducted in the presence of restraints. As the domain swapping process was carried out manually, no bonds were imposed at position 58 (in any subunit), nor for the two amino acids on either side of this residue (i.e. Ile56 and Val57, Gly59 and Val60), thereby permitting slow relaxation of the structure in the area closest to the domain exchange region that is located in the L1 loop (i.e. within the Ile56-Val60 segment). During the molecular dynamics simulations, the strength of these positional restraints was gradually reduced from an initial value of 20 kcal/mol to a final value of 0.3 kcal/mol.

During the second cycle of minimization, the MD simulations of the stab-1 HCC oligomers were carried out in *implicit* solvent (considering it to be an infinite medium, with specific properties related to water, such as the dielectric constant). The solvation model was described by the Born model as generalized in the Hawkins, Cramer, Truhlar approach [59, 60], using the parameters provided by Tsui& Case [61]. The lengths of the hydrogen bonds were maintained at constant values using the SHAKE algorithm, and for all of the oligomers the temperature was kept at 300K and was regulated by the Berendsen algorithm [62].

The total time for the simulation of each oligomer was 50 ns (i.e. 25 million steps), but the simulation procedure was split into several cycles. After each cycle, a model of the stab-1 HCC oligomer was generated and the positional restraints imposed on the Cα atoms were gradually reduced from 20 to 0.5 kcal/mol in the time range 0-35 ns, and to 0.3 kcal/mol in the time range of 35-50 ns. The MD simulations were continued until the energy of the system reached equilibrium.

The analysis of the oligomer models obtained during MD simulations involved (i) analysis of the trajectory (RMSD) performed using *ptraj* from the AMBER package, (ii) analysis of the hydrogen bonds using *cpptraj* from the AMBER package, (iii) analysis of the secondary structure using the STRIDE web server [44], and (iv) analysis of the potential energy and temperature using the process_mdout.perl script (AMBER) to extract the information from the MD output files. Visual representations of the structures of the oligomers were generated using PyMOL (*The PyMOL Molecular Graphics System, Version 1.5.0.4 Schrödinger, LLC*).

#### 4.5.3. Canonical molecular dynamics with use of scale-consistent UNitedRESidue (UNRES) coarse-grained force field

To determine the stability of the obtained models we performed canonical molecular dynamics using scale-consistent United RESidue (UNRES) coarse-grained force field as described previously [42]. We performed simulations at two temperatures: 277K (experimental temperature) and 300K (room temperature) with 48 trajectories for each system and each temperature. For both temperatures, we performed 6 million step simulations with 0.498 fs time steps that correspond to 3 ns of UNRES time which, after compensating for the speed due to coarse-graining, corresponds to ~ 3 μs of real-time [63, 64]. Langevin thermostat was used and the friction factor was scaled down by 100 to speed up the simulations. To prevent eventual reassociation of the dissociated multimers, the box size was set to 800 Å X 800 Å X 800 Å. Root-mean-square deviation (RMSD) measurements and TM-scores [43] were used to estimate the stability of the models.

#### 4.5.4. All-atom simulation with AMBER ff14SB force field

The all-atom simulations were performed with the AMBER ff14SB force field. The model protein was placed in a cuboid box in TIP3P explicit water. The size of the box was the size of the protein with an additional 10Å from each side for decamer Model 1 and 20 Å from each side for decamer Model 2 and the Dodecamer model, as the latter have larger sizes. The energy was minimized with a protocol that consisted of (i) restrained energy minimization with C*α*-distance restraints derived from the appropriate model and on the backbone dihedral angles from the regions of regular *α*-helical and *β*-sheet structure followed by (ii) a short restrained MD simulation with the same restraints. Subsequently, the short NPT simulation was performed and afterwards a 100 ns NVT production run was performed.

## Supplementary Materials

Supplementary materials can be found at www.mdpi.com/xxx/s1 and consist on ten figures: Figure S1. SAXS data recorded for the stab-1 HCC oligomers in solution; Figure S2. Modeling of the stab-1 HCC oligomers; Figure S3. The domain swapping scheme in the stab-1 HCC oligomers; Figure S4. Molecular dynamics simulations of the stab-1 HCC oligomers; Figure S5. UNitedRESidue (UNRES) coarse-grain simulations of the stab-1 HCC oligomers at 277K; Figure S6. UNitedRESidue (UNRES) coarse-grain simulations of the stab-1 HCC oligomers at 300K; Figure S7. UNitedRESidue (UNRES) coarse-grain simulation models of the stab-1 HCC oligomers at 277K; Figure S8. UNitedRESidue (UNRES) coarse-grain simulation models of the stab-1 HCC oligomers at 300K; Figure S9. RMSD as a function of time; Figure S10. Structure after 100ns coarse-grain simulation of Model 1 of decamer.

## Author Contributions

Conceptualization and methodology, M.C., J.R.K., M.K.; Investigation, M.C., A.K.S., S.R.-M., J.R.K., M.K.; Formal Analysis, M.C., A.K.S., S.R.-M., A.G., C.M.D., J.R.K., M.K.; Visualization, M.C., A.K.S. and M.K.; Writing - original draft, review & editing, M.C., A.K.S., S.R.-M., A.G., C.M.D., J.R.K., M.K. All authors have read and agreed to the published version of the manuscript.

