## Supplemental Information for "Structural characterization of covalently stabilized human cystatin C oligomers"

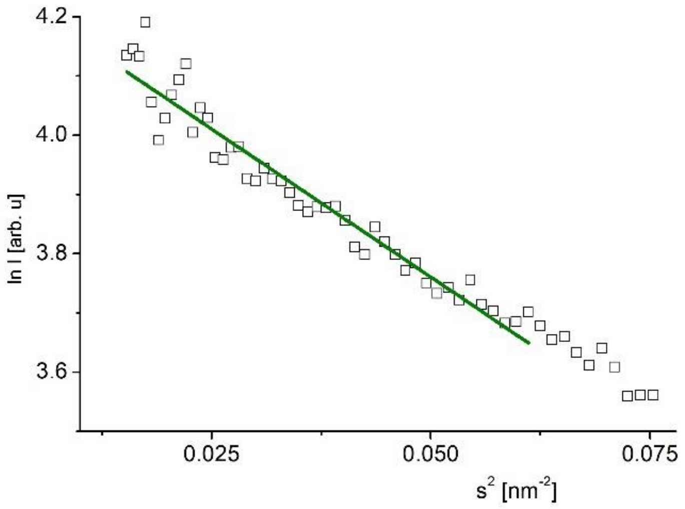


**Figure S1. SAXS data recorded for the stab-1 HCC oligomers in solution.** Guinier plot of SAXS data collected to define the size of the stab-1 HCC oligomers**.** The fit (green line) to the Guinier equation was performed within the SAXS data range of 0.124 to 0.247 $nm^{-1}$, where $sR_g < 1.3$ as the particles are assumed to be spherical, resulting in a value of $R_g$ of 5.28 ± 0.3 nm.

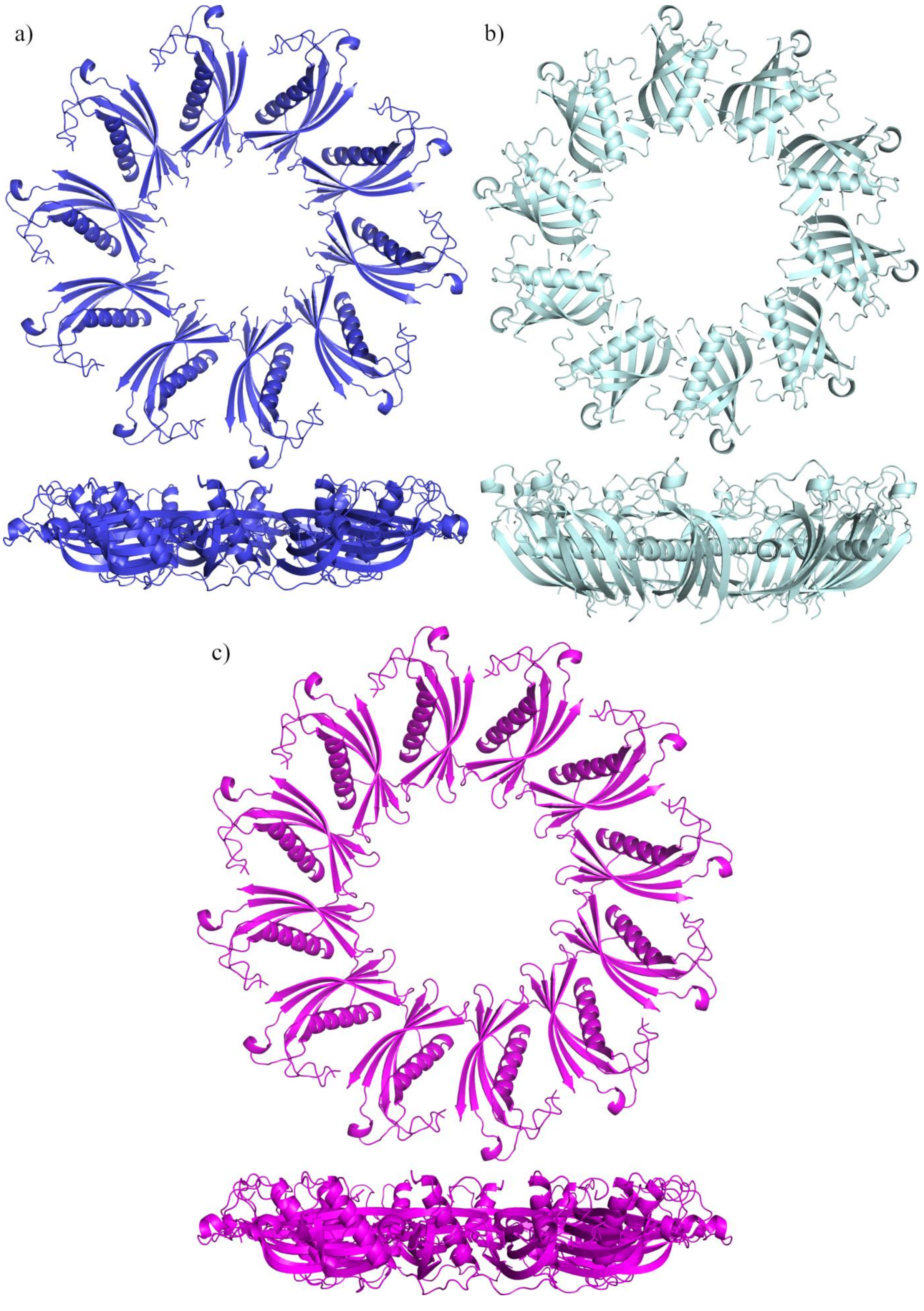


**Figure S2. Modelling of the stab-1 HCC oligomers.**

Preliminary models selected for MD simulations from set of solutions obtained from SymmDock: **a)** Model 1 of decamer; **b)** Model 2 of decamer; **c)** Model of dodecamer.

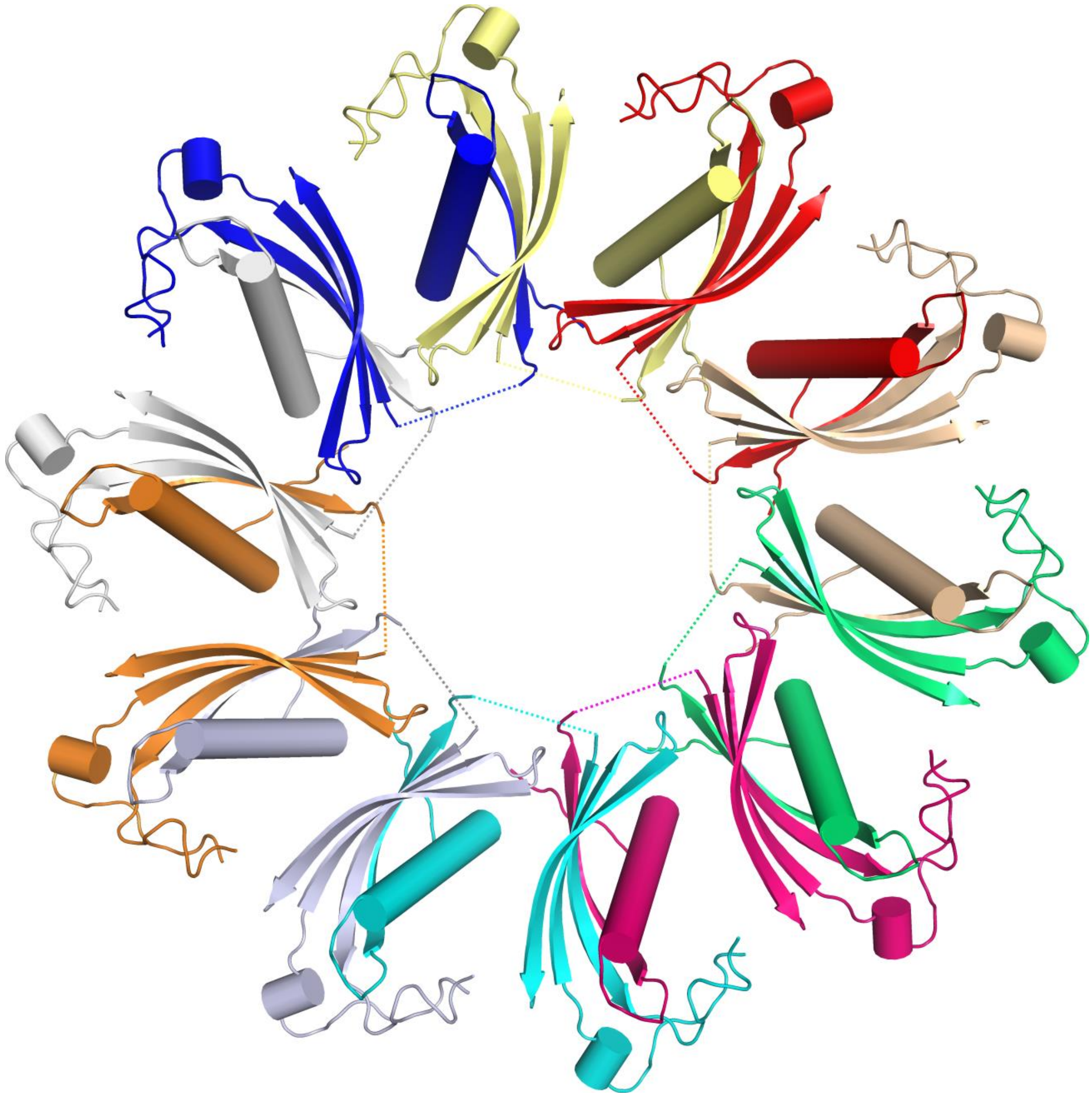

**Figure S3. The domain swapping scheme in the stab-1 HCC oligomers.**
The domain exchange between adjacent subunits within the stab-1 HCC oligomer from the Model 1 decamer. The colours indicate the individual stab-1 HCC subunits. To implement the domain swap, the bond between residues 58 and59 were manually broken and the atomic positions of the N-terminal fragment of the polypeptide chain (residues up to 58) was transferred from the first stab-1 HCC subunit to the next one, with the rest of the chain remaining unaltered. The dashed lines indicated the broken bond between residues 58 and 59.

**a)**

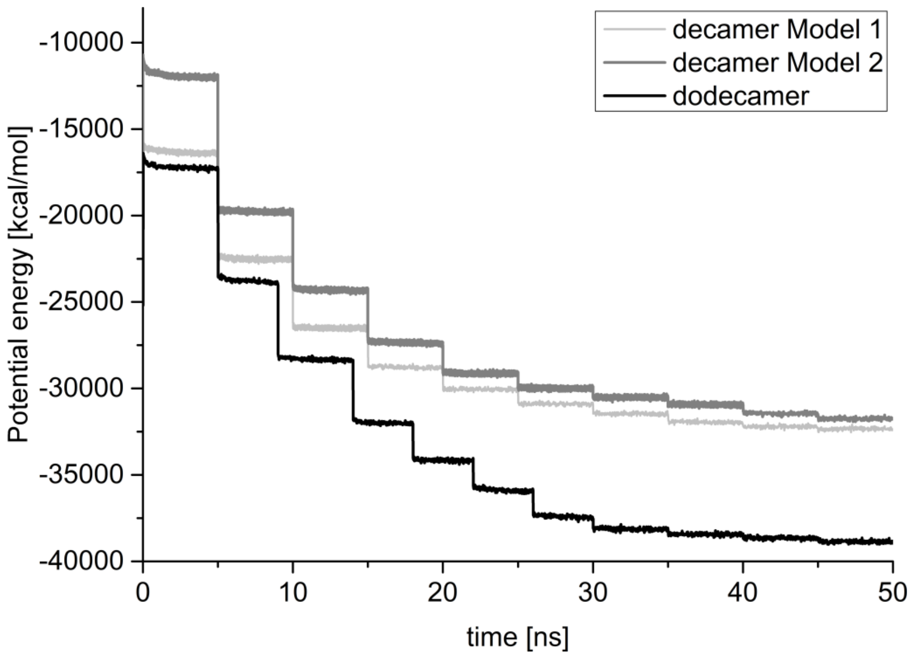


**b)**

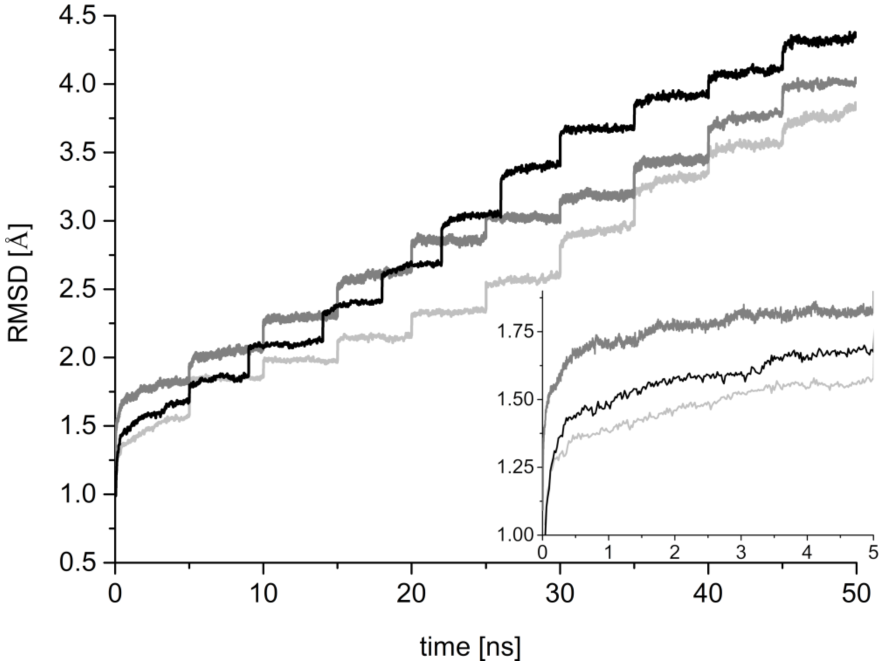


**Figure S4. Molecular dynamics simulations of the stab-1 HCC oligomers.**
**(a)** The potential energy ($E_{pot}$) of the MD system as a function of simulation time for HCC oligomers models. **(b)** The root-mean-square deviation (RMSD) of all $C_\alpha$ atoms calculated by reference to the initial structure taken as a function of the simulation time. (Inset) An expanded view of first 5 ns cycle (light gray - Model 1 of decamer; gray - Model 2 of decamer, dark gray – dodecamer).

**a)**

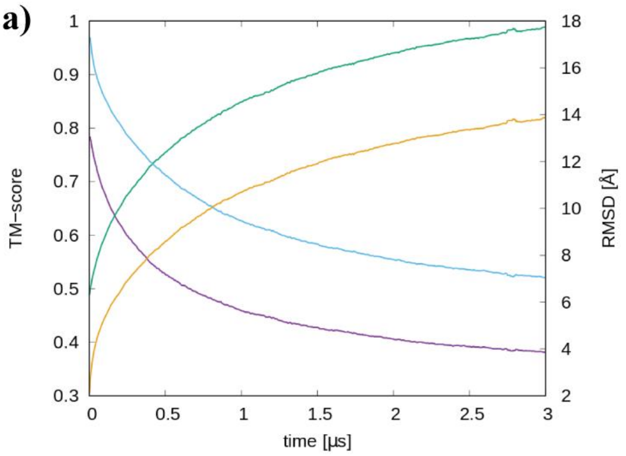


**b)**

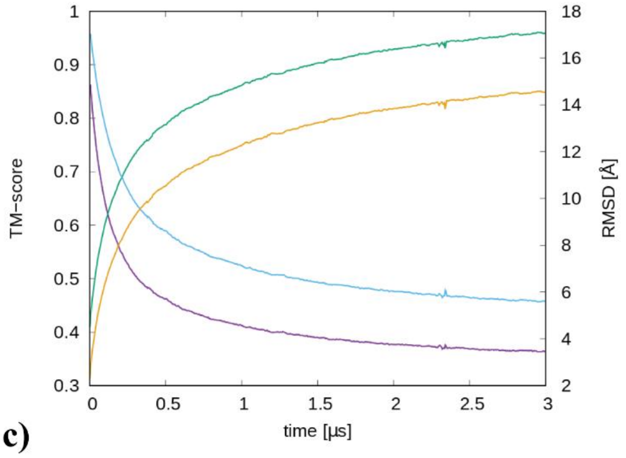


**c)**

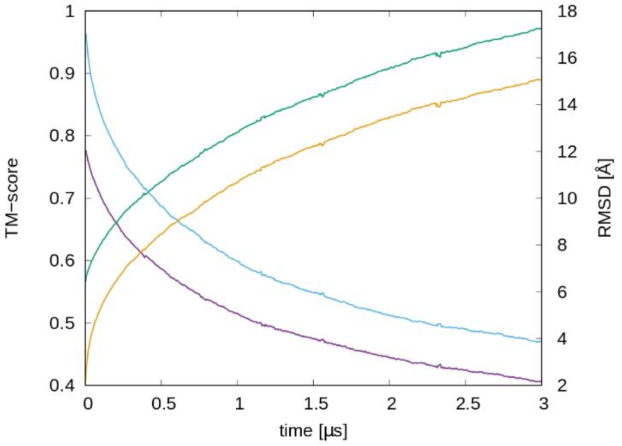


**Figure S5. UNitedRESidue (UNRES) coarse-grain simulations of the stab-1 HCC oligomers at 277K.**

The root-mean-square deviation (RMSD) and TM-scores for **a)** dodecamer, **b)** decamer Model 1, and **c)** decamer Model 2 at 277K over all trajectories. The blue and violet lines represent the TM-score plots for the UNRES minimized and original (reference) model, respectively. The yellow and green lines represent the RMSD plots for the UNRES minimized and original (reference) model, respectively.

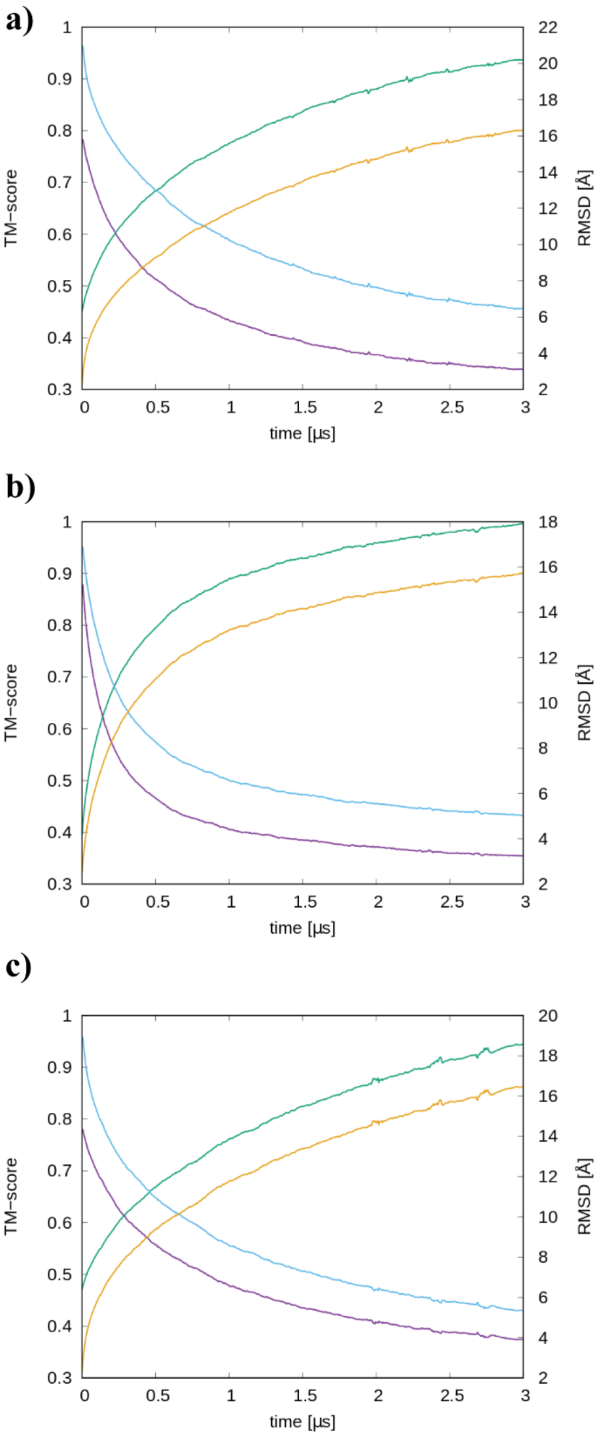


**Figure S6. UNitedRESidue (UNRES) coarse-grain simulations of the stab-1 HCC oligomers at 300K.**

The root-mean-square deviation (RMSD) and TM-scores for **a)** dodecamer, **b)** decamer Model 1, and **c)** decamer Model 2 at 300K over all trajectories. The blue and violet lines represent the TM-score plots for the UNRES minimized and original (reference) model, respectively. The yellow and green lines represent the RMSD plots for the UNRES minimized and original (reference) model, respectively.

**a)**

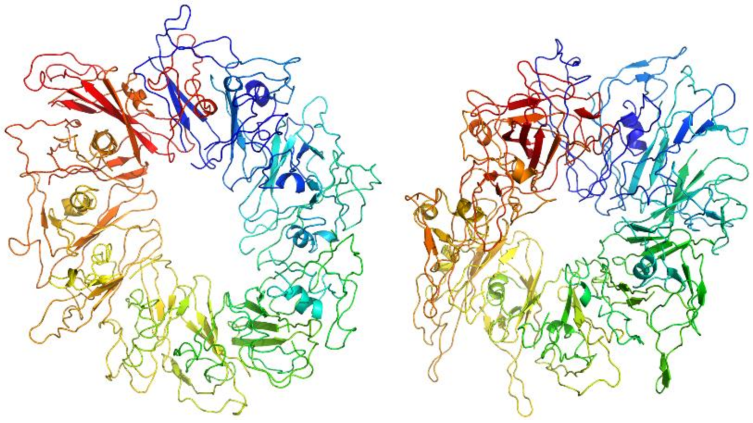

**b)**

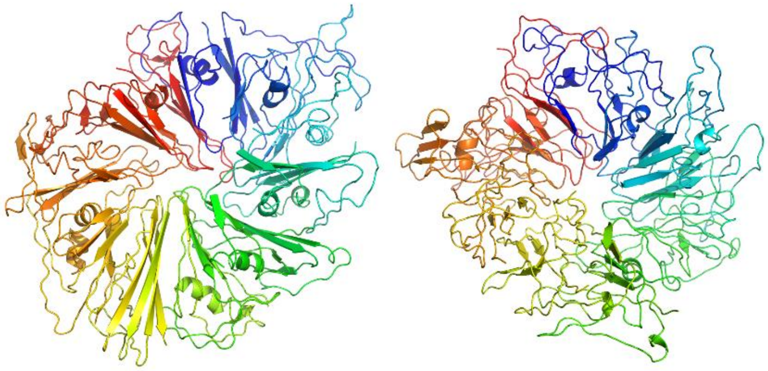

**c)**

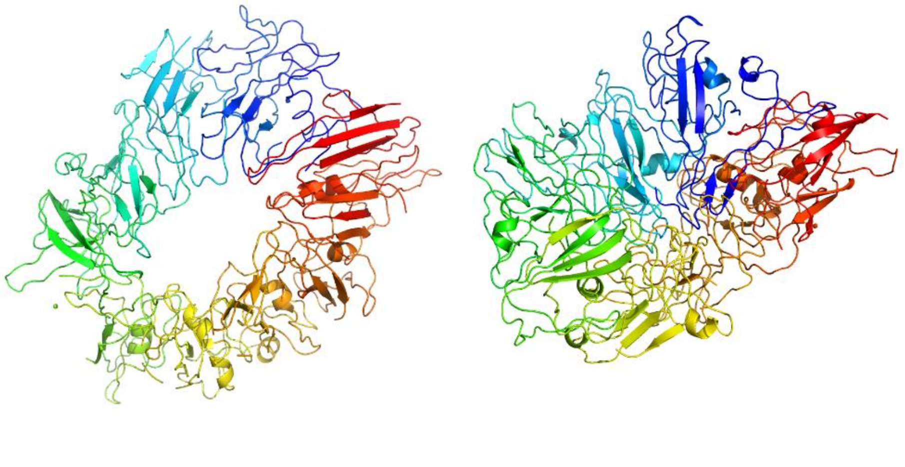

**Figure S7. UNitedRESidue (UNRES) coarse-grain simulation models of the stab-1 HCC oligomers at 277K**

Representative models after long-term UNRES coarse-grain simulations **a)** dodecamer, **b)** decamer Model 1 and **c)** decamer Model 2 at 277K.

a)

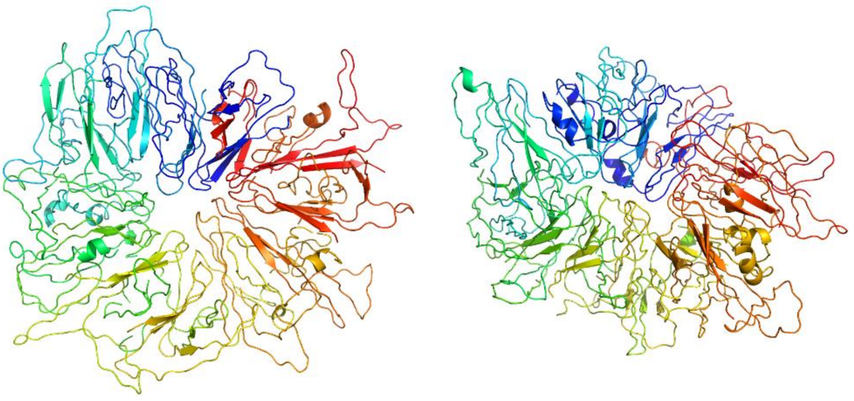

b)

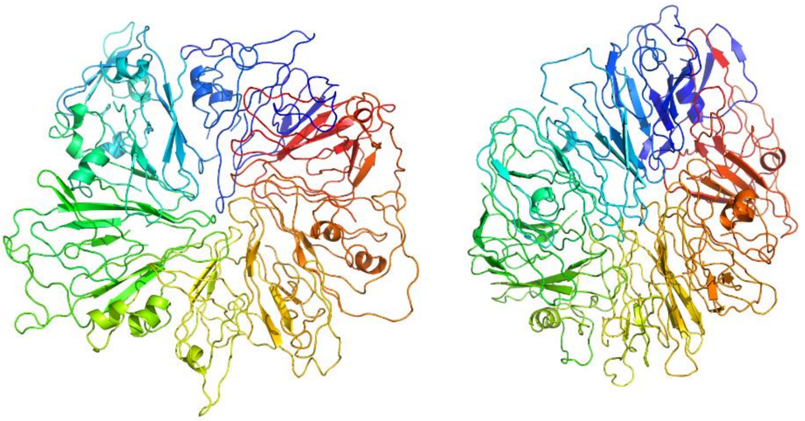

c)

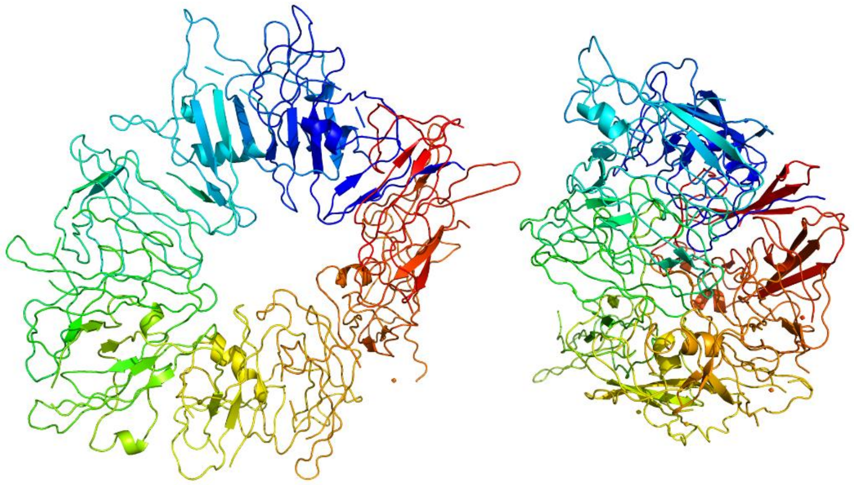

**Figure S8. UNitedRESidue (UNRES) coarse-grain simulation models of the stab-1 HCC oligomers at 300K**

Representative models after long-term UNRES coarse-grain simulations **a)** dodecamer, **b)** decamer Model 1 and **c)** decamer Model 2 at 300K.

**a)**

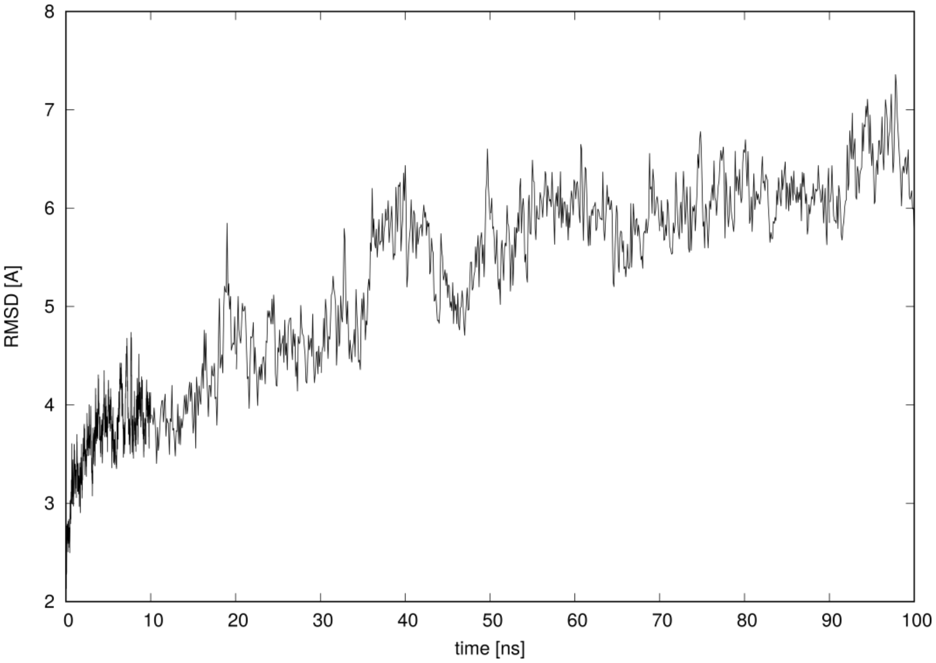


**b)**

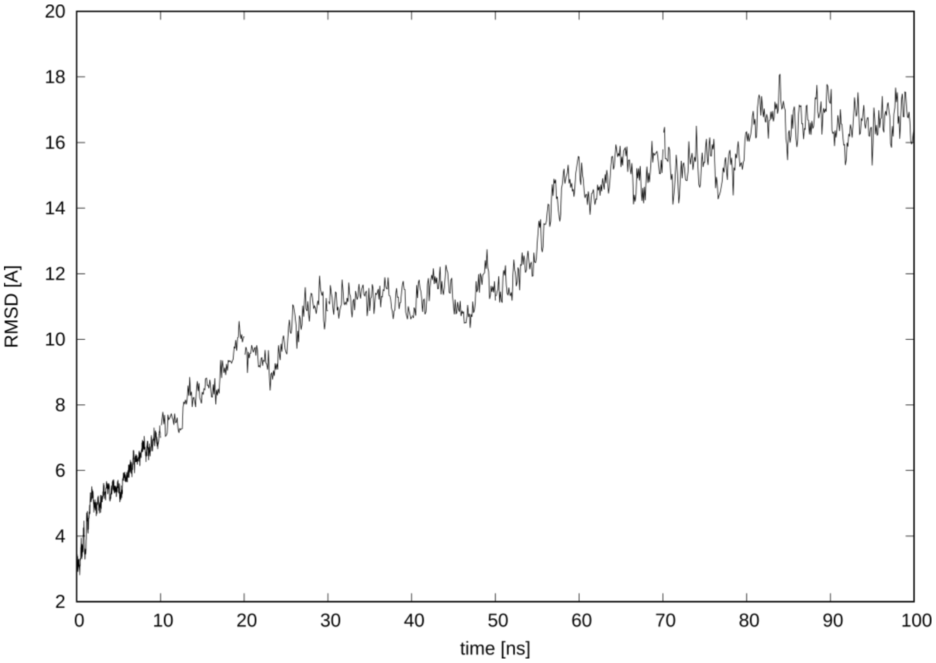


**Figure S9 RMSD as a function of time in UNitedRESidue (UNRES) coarse-grain simulations for a) Model 1 decamer and b) dodecamer.**

**a)**

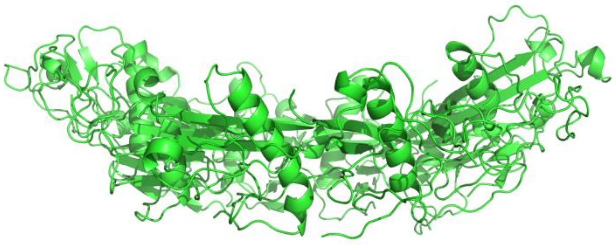

**b)**

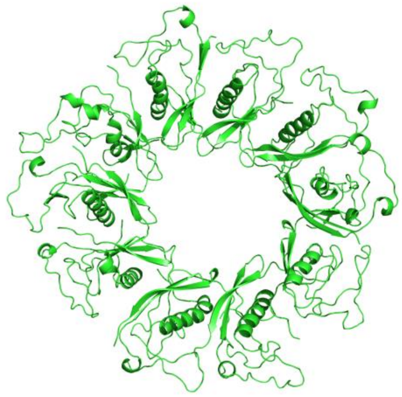

**c)**

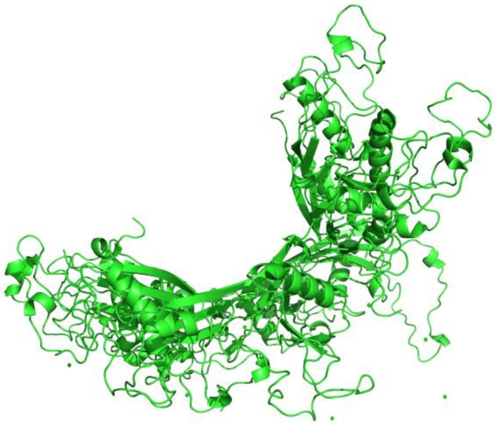

**d)**

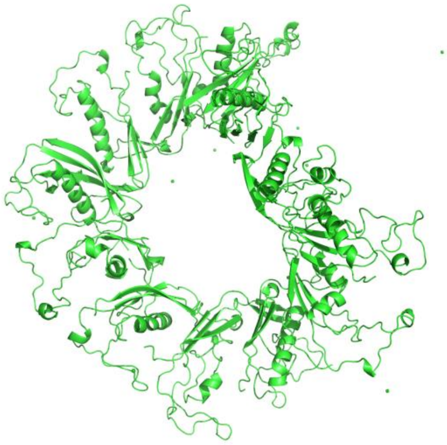

**Figure S10 Structure after 100ns coarse-grain simulation** of Model 1 of decamer.

**a)** side view and **b)** top view compared with structure obtained after 100ns simulation of dodecamer **c)** side view and **d)** top view.
